# Eelgrass leaf surface microbiomes are locally variable and highly correlated with epibiotic eukaryotes

**DOI:** 10.1101/111559

**Authors:** Mia M. Bengtsson, Anton Bühler, Anne Brauer, Sven Dahlke, Hendrik Schubert, Irmgard Blindow

**Affiliations:** Institute of Microbiology, University of Greifswald, Greifswald, Germany; Institut fur Biowissenschaften, University of Rostock, Rostock, Germany; Biological Station of Hiddensee, University of Greifswald, Kloster, Germany

## Abstract

Eelgrass *(Zostera marina)* is a marine foundation species essential for coastal ecosystem services around the northern hemisphere. Like all macroscopic organisms, it possesses a microbiome which may play critical roles in modulating the interaction of eelgrass with its environment. For example, its leaf surface microbiome could inhibit or attract eukaryotic epibionts which may overgrow the eelgrass leading to reduced primary productivity and subsequent eelgrass meadow decline. We used amplicon sequencing of the 16S and 18S rRNA genes of prokaryotes and eukaryotes to assess the leaf surface microbiome (prokaryotes) as well as eukaryotic epibionts in-and outside lagoons on the German Baltic Sea coast. Bacterial microbiomes varied substantially both between sites inside lagoons and between open coastal and lagoon sites. Water depth, leaf area and biofilm chlorophyll *a* concentration explained a large amount of variation in both bacterial and eukaryotic community composition. Communities of bacterial and eukaryotic epibionts were highly correlated, and network analysis revealed disproportionate co-occurrence between a limited number of eukaryotic taxa and several bacterial taxa. This suggests that eelgrass leaf surface biofilms are a mosaic of the microbiomes of several eukaryotes, in addition to that of the eelgrass itself, and underlines that eukaryotic microbial diversity should be taken into account in order to explain microbiome assembly and dynamics in aquatic environments.

## INTRODUCTION

Seagrasses are aquatic flowering plants that form underwater meadows critically important for coastal ecosystems around the world. Seagrass meadows are nursing grounds for juvenile fish which can hide and forage between the seagrass leaves, sediments are stabilized by the seagrass roots and the biomass of seagrass and associated organisms sequester carbon with implications for climate change mitigation (Fourqurean et al., 2012). The seagrass eelgrass *(Zostera marina*) is an important foundation species along soft-bottom coasts in the northern hemisphere. Eelgrass meadows are biodiversity hotspots, providing a home to a myriad of invertebrates, algae, fish and microorganisms. Due to its ability to tolerate low and fluctuating salinity levels, eelgrass is common in estuaries as well as in the biggest brackish water habitat in the world, the Baltic Sea. However, significant declines in the depth limit and areal cover of eelgrass meadows in the Baltic Sea have been observed since several decades (Boström et al., 2014). This echoes the distressing situation for seagrass ecosystems around the world, which experience threats by human activities and global change (Orth et al., 2006).

The mechanisms that are responsible for eelgrass meadow decline appear to depend on many different factors. Historically, outbreaks of the pathogenic protist *Labyrinthula zosterae* (“wasting disease”) has caused catastrophic die-off of eelgrass (Muehlstein et al., 1991), although its importance as a pathogen under current conditions in Europe seems limited (Brakel et al., 2014). Instead, factors such as water clarity, eutrophication, grazing pressure, and interspecific competition have been identified as culprits for eelgrass growth (Baden et al., 2010;Duffy et al., 2015).

Especially important is the competition between eelgrass and algae that grow within meadows, often as epiphytes on the eelgrass itself. In some cases, eelgrass becomes extensively covered with a mixture of algae, bacteria and sessile animals, which shade it and inhibit transport of solutes (Brodersen et al., 2015) and may over time cause eelgrass meadows to degrade. Recent research suggests that relative success of eelgrass and associated algae is determined by complex interactions between biotic processes such as grazing on both algae and eelgrass by invertebrates, and environmental factors such as nutrient concentrations and temperature (Alsterberg et al., 2013; Eklo &#x00F6;f et al., 2012). However, these mechanisms are not well understood and their complexity necessitates a holistic ecosystem approach taking into account several organism groups that inhabit seagrass meadows and their interactions to be resolved (Boström et al., 2014;Maxwell et al., 2016).

An overlooked group of organisms in eelgrass meadows is the bacteria, which cover eelgrass leaves and roots forming biofilms, are responsible for degradation of eelgrass detritus and are also associated to all other organisms in eelgrass beds including algae and other epiphytes. Bacteria are the first colonizers on new seagrass leaves and thus initiate a successional process that may end with severe epiphytic overgrowth. Bacteria are also an important food source for microbial eukaryotic and invertebrate grazers and may therefore support a substantial part of the foodweb found within eelgrass meadows. Research on eelgrass leaf bacterial communities is very limited, but early work has determined that bacterial abundance and productivity on eelgrass leaves vary during the year, with a peak in early autumn (Tornblom and Sondergaard, 1999), and that eelgrass leaves have an associated bacterial community (the “microbiome”), with only some compositional overlap with other aquatic macrophytes (Crump and Koch, 2008).

There is currently little known about what factors influence eelgrass microbiomes, or what functions the bacteria have on eelgrass leaves. Bacteria are likely to play a fundamental role in the competition of eelgrass and epibionts such as algae. For example, certain bacteria may inhibit the attachment of algal spores while others may promote further colonization (Celdra &#x00E1; n et al., 2012;Mieszkin et al., 2013). Conversely, epibiotic eukaryotes including algae may in turn shape the bacterial communities on eelgrass leaves by release of bacterial attractants or deterrents (Steinberg and de Nys, 2002) or through selective grazing (Huws et al., 2005), for example. Biotic and abiotic environmental factors as well as host-related factors such as eelgrass productivity and genotype are also likely to shape the eelgrass microbiome, and thereby modulate its function.

In this study, we aimed to obtain a first view onto eelgrass leaf microbiomes by investigating the community composition of prokaryotic and eukaryotic epibiotic organisms in relation to abiotic and biotic environmental variables. We sampled eelgrass in semi-sheltered lagoons and along exposed open shorelines around the Island of Hiddensee on the eastern German Baltic Sea coast and used high-throughput Illumina amplicon sequencing of the 16S and 18S rRNA genes. We hypothesized that (1) both bacterial and eukaryotic communities would vary substantially between lagoon and open coast sites due to different abiotic conditions and that (2) bacterial microbiome composition would be influenced by the composition of epibiotic eukaryotes.

## MATERIALS & METHODS

### Study area

We chose to perform our survey in the area around the island of Hiddensee on the German Baltic Sea coast as it offers variable yet locally representative environments colonized by eelgrass meadows. The area is characterized by semi-enclosed, shallow lagoons (German: “Bodden”) which differ from the open coast with respect to depth distribution, salinity and nutrient concentrations. The surveyed lagoons (the Vitter Bodden and Schaproder Bodden) feature somewhat lower average salinity (8.8 ± 1.0 PSU, range 6.5 - 13.4 PSU) compared to the open coast (Libben bay, 9.4 ±1.6 PSU, range 6.8 - 15.9 PSU). Nutrient levels are elevated in lagoons (total N: 38.2 ±12.3, total P: 1.2 ±0.62 q &#x03BC;mol l^−1^) compared to open coast waters (total N 19.9 ±4.3, total P: 0.91 ±0.4 q &#x03BC;mol l^−1^) mainly due to agricultural runoff and other human activities (Schiewer, 2008). Salinity and nutrient values are yearly averages from monitoring data 2005 –2014 (Landesamt für Umwelt, Naturschutz und Geologie Mecklenburg-Vorpommern, unpublished data).

#### Sampling

*Zostera marina* shoots were sampled via scuba diving at 7 sites around the island of Hiddensee (Fig. 1). At every site, 5 replicate shoots (above-ground parts) were collected within an area of approximately 1 m^2^. Shoots were kept cool and in clean plastic bags until sample processing which was carried out within a few hours. Environmental variables such as water depth and parameters relating to the surrounding macrophyte vegetation (eelgrass and co-occurring macroalgae and aquatic plants) were determined as part of a simultaneous macrophyte inventory (Bu&#x00FC; hler, 2016).

**Figure 1:**
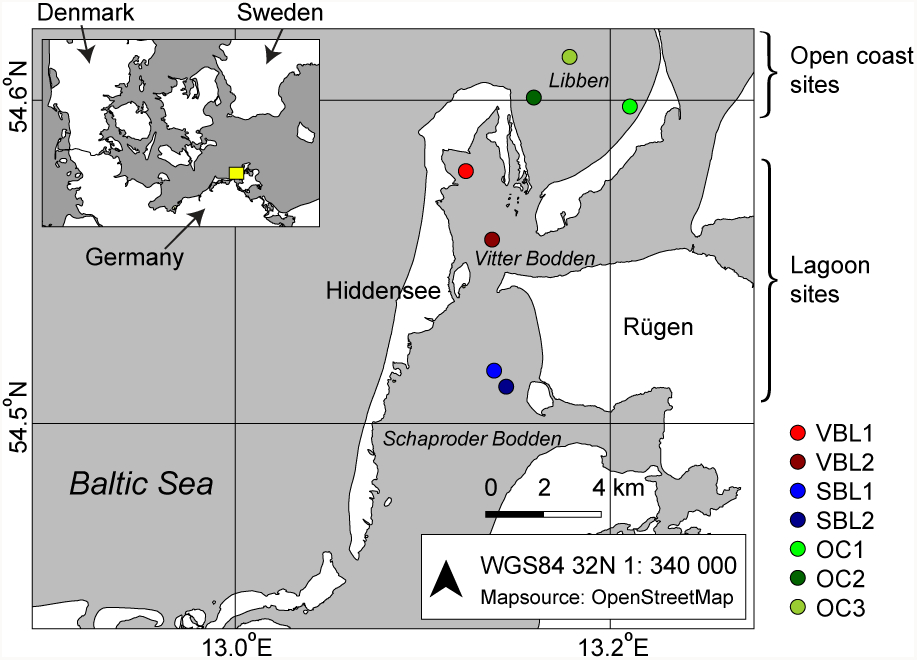
Location of sampling sites around the island of Hiddensee on the German Baltic Sea coast. The inset shows the geographic placement of the area.

#### Leaf biofilm sample processing

The sampled eelgrass shoots were rinsed with sterile filtered seawater and handled under sterile conditions. Leaf areas with heavy macroscopic epibiosis and visibly degrading tissue were removed before biofilm sampling. Epibiotic biofilm from the entire leaf surface of each shoot was scraped off using sterile cotton swabs. The scraped off material was suspended in sterile-filtered seawater (10 ml) and kept cool. The suspensions were aliquoted into separate tubes and centrifuged (10000 x g, 10 min., 4°C) to pellet biofilm material. Pellets were frozen at −20°C until DNA and chlorophyll *a* extraction.

Chlorophyll *a* was extracted overnight in acetone at +4° C and absorbance at 665 nm and 750 nm was measured on a spectrophotometer (Genesys 20, Thermo Fisher Scientific). Biofilm chlorophyll *a* content was calculated as p&#x03BC;g chlorophyll *a* cm^−2^ of biofilm.

#### DNA extraction and Illumina amplicon sequencing

DNA was extracted from biofilm pellets using the MoBio PowerSoil kit for DNA (MOBIO Laboratories). Mechanical lysis was achieved by bead beating in a FastPrep 24 5G (MP Biomedicals). Extracted DNA was amplified with primer pairs targeting the V4 region of the 16S rRNA (515f: 5’- GTGYCAGCMGCCGCGGTAA-3 ’, 806r: 5’-GGACTACNVGGGTWTCTAAT-3’ (Walters et al., 2015) and the V7 region of the 18S rRNA (F-1183mod: 5’- AATTTGACTCAACRCGGG-3’, R-1443mod: 5’- GRGCATCACAGACCTG-3’ (Ray et al., 2016)) coupled to custom adaptor-barcode constructs. PCR amplification and Illumina MiSeq library preparation and sequencing (V3 chemistry) was carried out by LGC Genomics in Berlin.

#### Sequence processing

Sequences clipped from adaptor and primer sequence remains were processed using the DADA2 package in R (version 1.2.0 Callahan et al., 2016; R Development Core Team, 2017). Briefly, forward and reverse Illumina reads were truncated to 200 bp, filtered (maxEE=2, truncQ=2), dereplicated and error rates were estimated using the maximum possible error estimate from the data as an initial guess. Sample sequences were inferred, and paired forward and reverse reads were merged. Chimeric sequences were removed using the removeBimeraDenovo function. The resulting unique sequence variants (analogous to operational taxonomic units) were used to construct a table containing relative abundances of sequence variants across all samples. Sequence variants were taxonomically classified using a lowest common ancestor approach (Lanze&#x00E9; n et al., 2012) based on the Silva database (Pruesse et al., 2007).

#### Statistical analyses

Multivariate statistical analyses were carried out using functions in the vegan package (version 2.4-1,Oksanen et al. (2016)) in R (version 3.3.1, R Development Core Team, 2017). To visualize similarities in sequence variant composition (community composition) between sampling sites, non-metric multidimensional scaling (vegan function metaMDS) was performed on Hellinger-transformed sequence variant counts using Bray-Curtis distance. To explain the variation in community composition in response to the environment, abiotic and biotic variables were selected based on their explanatory power in PERMANOVA tests (vegan function adonis, variables were transformed (log_e_) to achieve normal-distribution if needed). A final model including the two most relevant environmental variables was selected and distance-based redundancy analysis ordination (vegan function capscale, Bray-Curtis distance) constrained to these environmental variables was used to visualize variation.

Sequence variant richness was calculated by rarefying read counts to the lowest number of reads in a sample of each dataset (16S: 5861, 18S: 25780). Evenness was calculated as J = H/logS, where H is the Shannon diversity index and S is rarefied richness (Pielou, 1977). To identify sequence variants that were differently abundant in lagoon sites and open coast sites, differential abundance analysis as implemented in the DEseq2 R package was used (Love et al., 2014).

A partial mantel test was used to assess correlation between bacterial and eukaryotic community composition while correcting for environmental variables (vegan function mantel.partial, Bray-Curtis distance for community data, euclidean distance for environmental variables,Legendre and Legendre (2012)). To analyze pairwise correlations between 16S and 18S sequence variants, co-occurrence networks were calculated in R using Pearson correlation. P-values were adjusted for multiple testing using the Benjamini-Hochberg method (Benjamini and Hochberg, 1995). Only correlations with adjusted p-values < 0.01 were treated as significant and included in the network. The network was plotted and edited for clarity (symbols, colors etc.) in Cytoscape version 3.3.0 (Shannon et al., 2003).

### RESULTS

Due to the different depth distributions of eelgrass on open coasts and in lagoons (Buehler, 2016), all open coast sites where eelgrass was found were deeper than lagoon sites. Eelgrass biomass (leaf surface area and dry weight) varied widely among sampling sites (Table 1). Leaf surface biofilms were visually different in color and density (results not shown) and biofilm Chlorophyll *a* content varied substantially among sampling sites (Table 1).

**Table 1:** An overview of the sampling sites and key abiotic and biotic parameters. Leaf area and chlorophyll *a* values represent the mean of 5 replicates ± 1 standard deviation. Water depth is the depth at which eelgrass was collected.

| Site | Water body | Latitude<br>Longitude | Water<br>depth (m) | Eelgrass leaf<br>area ($\text{cm}^2$) | Biofilm chlorophyll<br>*a* ($\mu\text{g cm}^{-2}$) | Eelgrass dry<br>weight ($\text{g m}^{-2}$) |
| --- | --- | --- | --- | --- | --- | --- |
| VBL1 | Lagoon | 54.57827<br>13.12357 | 2.5 | 391.1<br>$\pm 130.7$ | 0.053<br>$\pm 0.018$ | 536.95 |
| VBL2 | Lagoon | 54.55705<br>13.13677 | 3.0 | 251.6<br>$\pm 42.3$ | 0.011<br>$\pm 0.003$ | 230.54 |
| SBL1 | Lagoon | 54.51641<br>13.13775 | 3.4 | 292.7<br>$\pm 50.5$ | 0.021<br>$\pm 0.002$ | 107.70 |
| SBL2 | Lagoon | 54.51144<br>13.14459 | 1.9 | 67.0<br>$\pm 16.1$ | 0.052<br>$\pm 0.016$ | 5.00 |
| OC1 | Open coast | 54.59808<br>13.21117 | 5.0 | 151.2<br>$\pm 49.7$ | 0.017<br>$\pm 0.002$ | 72.04 |
| OC2 | Open coast | 54.60116<br>13.15891 | 3.9 | 140.4<br>$\pm 30.8$ | 0.025<br>$\pm 0.005$ | 17.14 |
| OC3 | Open coast | 54.61305<br>13.17829 | 7.1 | 139.7<br>$\pm 25.7$ | 0.037<br>$\pm 0.006$ | 47.30 |

Illumina amplicon sequencing of 16S and 18S rRNA fragments resulted in 1.0 million 16S reads and 1.9 million 18S reads with an average of 29564 16S reads (min = 5861, max = 67456) and 56886 reads per sample (min = 25780 max = 126938). One sample from site SBL was excluded from all analyses because of low 16S read count (861 reads). In total, 4409 prokaryotic and 1946 eukaryotic sequence variants (analogous to operational taxonomic units) were identified. Among prokaryotes, *Bacteria* were overwhelmingly abundant over *Archaea* (99.2 *%* of sequence variants and 99.9 % of reads). Due to this, 16S sequence variants are referred to as “*Bacterid*’ from here on.

Bacterial community composition varied substantially among sites (PERMANOVA R^2^ = 81.6, p < 0.001), and the main variation was between lagoon sites and open coast sites (PERMANOVA R^2^ = 42.4, p < 0.001, Fig. 2a). Eukaryotic community composition also varied among individual sites (PERMANOVA R^2^ = 82.7, p < 0.001) and varied strongly between lagoon and open coast sites (PERMANOVA R^2^ = 47.8, p < 0.001, Fig. 2b). Water depth, eelgrass shoot leaf area and biofilm chlorophyll *a* content were the strongest environmental predictors of community variation in both bacteria and eukaryotes, with depth explaining about 30 %, leaf area 11 % and chlorophyll *a* 10 % of total variation (PERMANOVA p < 0.001). There was also a significant interaction term between depth and chlorophyll *a* content for both bacterial and eukaryote communities (PERMANOVA, p < 0.01), indicating that chlorophyll *a* affects communities differently depending on depth, or vice-versa. The variation explained by depth and chlorophyll *a* content is illustrated in Fig. 2c–d.

**Figure 2:**
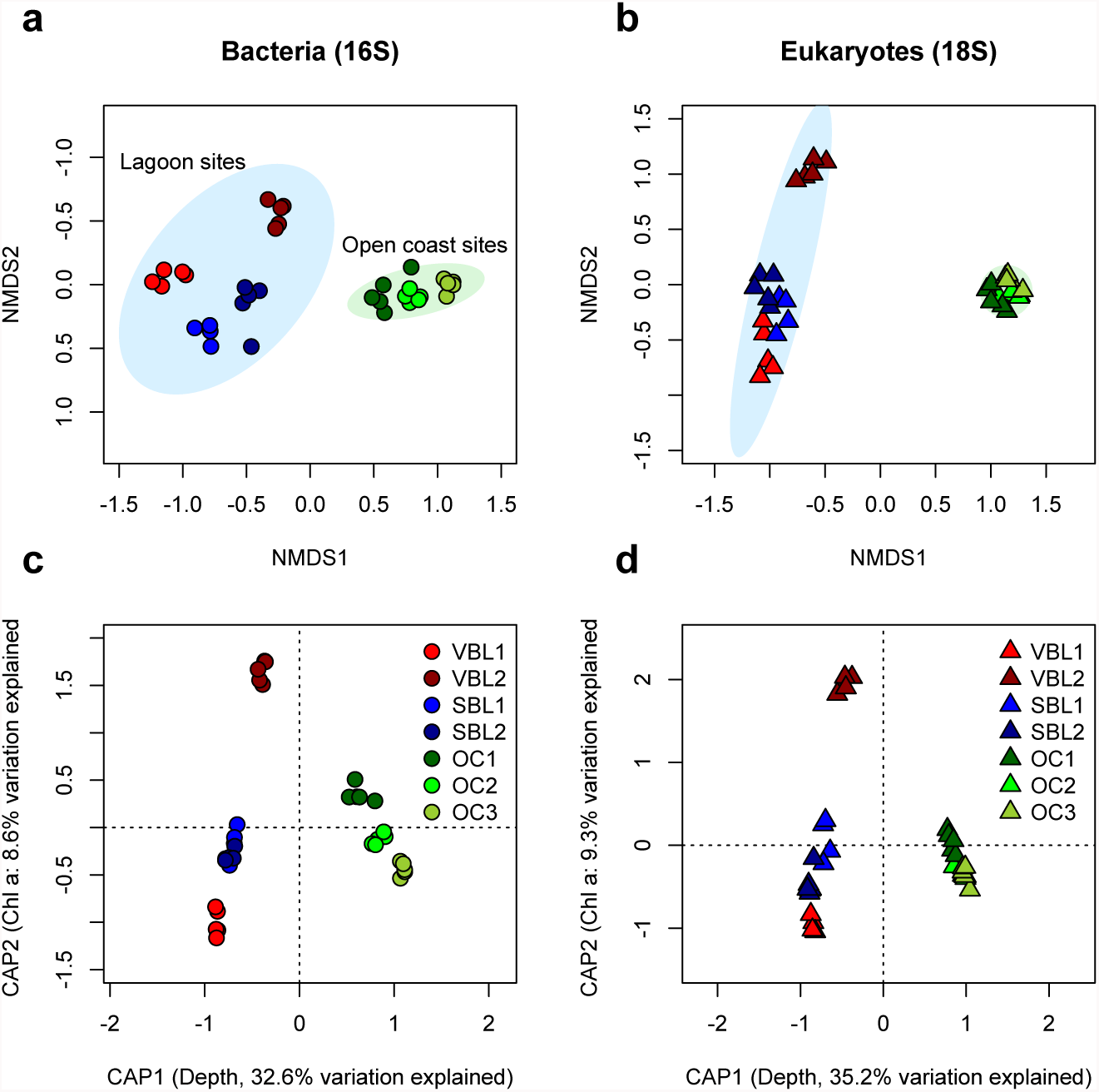
Ordinations based on 16S rRNA and 18S rRNA sequence variant relative abundances illustrate the variability in bacterial and eukaryotic community composition between sampling sites. In a) and b), non-metric multidimensional scaling (nMDS) display the distinct clustering of lagoon (blue ellipse) and open coast sites (green ellipse). Ellipses represent 95% confidence intervals of site position. In c) and d), distance-based RDA (Bray-Curtis distance metric used) constrained to water depth and leaf surface biofilm chlorophyll *a* content show the variability that can be explained by these two environmental variables.

Richness of bacterial sequence variants was positively correlated with eelgrass dry weight at the sampling site (R^2^ = 0.69, p < 0.001, Fig. 3a) as well as with eelgrass leaf area, although somewhat more weakly (R^2^ = 0.29, p < 0.001). Eukaryotic sequence variant richness did not correlate with eelgrass dry weight or other environmental gradients, yet was significantly higher in lagoon sites compared to open coast sites (ANOVA F = 208.6, p < 0.001, Fig. 3b). Richness and evenness were correlated for both bacteria (R^2^ = 0.34, p < 0.001) and eukaryotes (R^2^ = 0.84, p < 0.001).

**Figure 3:**
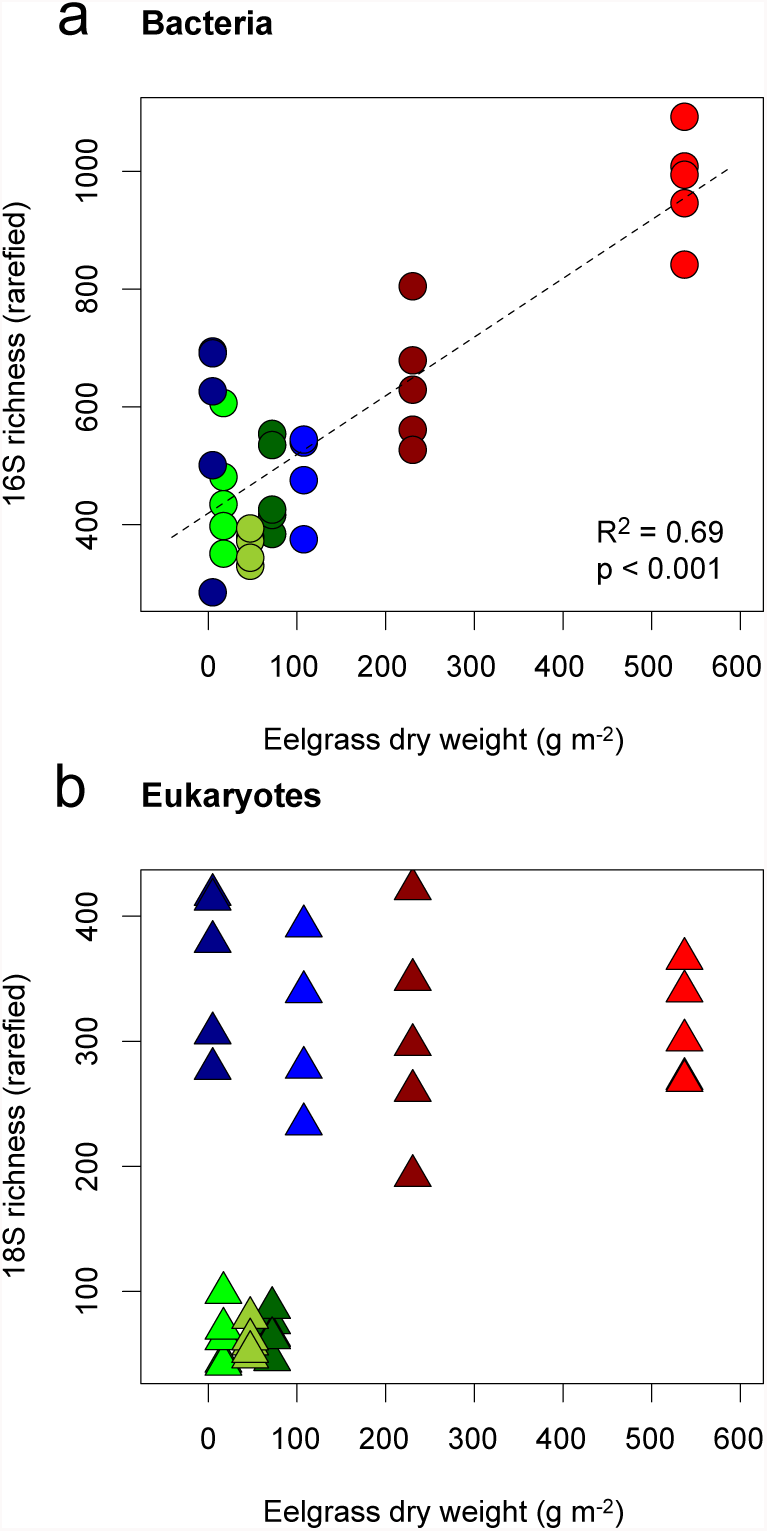
Richness of 16S (a) and 18S (b) sequence variants in relation to the biomass (dry weight) of *Zostera marina* at all sampling sites. Symbols and colors are the same as in Fig. 1.

A differential abundance analysis revealed sequence variants that differed significantly (adjusted p < 0.01) in relative abundance between lagoon and outer coast sites (Fig. 4). On a broad taxonomical level, several bacterial sequence variants from *Cyanobacteria, Planctomycetes, Gemmatimonadetes* and *Acidobacteria* were overrepresented in lagoon sites (Fig. 4a) whereas sequence variants from almost all eukaryotic taxonomical groups were overrepresented in lagoon sites (Fig. 4b). However, one highly abundant individual sequence variant, classified as *Hydrozoa (Metazoa: Cnidaria)* was significantly overrepresented in open coast sites, while several other metazoan (e.g. classified as *Dorvillea* and *Rotifera*) and diatom sequence variants (*Cocconeidaceae*) were overrepresented in lagoon sites. 14 bacterial and 3 eukaryotic sequence variants were detected in all samples, and were defined as the “core microbiome” of eelgrass leaf biofilms for the purpose of this study (Table 2).

**Figure 4:**
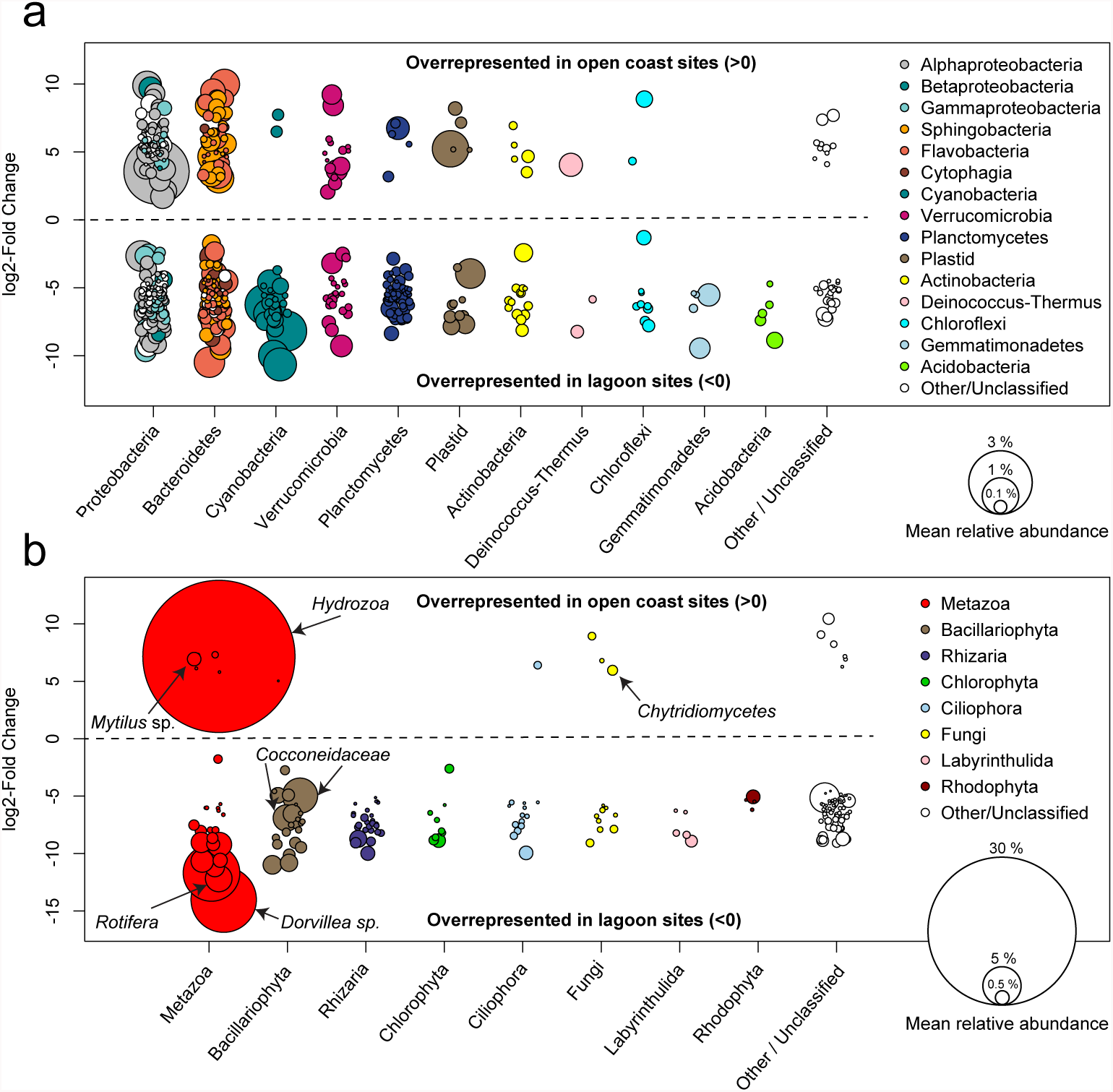
Differential abundance analysis of 16S (a) and 18S (b) sequence variants comparing open coast and lagoon sites. Positive log2 fold-change values indicate a significantly (adjusted p < 0.01) higher abundance in open coast sites, while negative values indicate significantly higher abundances in lagoon sites. The area of each circle representing an individual sequence variant is proportional to the relative abundance of that sequence variant across all samples. Prominent eukaryotic sequence variants are labeled with more detailed taxonomic classification.

**Table 2:** Mean relative abundance (across all samples) and taxonomic classifications of bacterial and eukaryotic sequence variants detected in all samples, i.e. the “core microbiome” of eelgrass leaf biofilms in the present study.

| Sequence variant id | Mean abundance (%) | Domain | Level 1 | Level 2 | Level 3 | Level 4 |
| --- | --- | --- | --- | --- | --- | --- |
| bac4 | 3.21 | Bacteria | Proteobacteria | Alphaproteobacteria | Rhodobacterales | Rhodobacteraceae |
| bac7 | 2.73 | Bacteria | Proteobacteria | Betaproteobacteria | Methylophilales | Methylophilaceae |
| bac9 | 2.52 | Bacteria | Proteobacteria | Alphaproteobacteria | Rhodobacterales | Rhodobacteraceae |
| bac17 | 1.33 | Bacteria | Proteobacteria | Alphaproteobacteria | Rhodobacterales | Rhodobacteraceae |
| bac28 | 1.27 | Bacteria | Proteobacteria | Alphaproteobacteria | Rhodobacterales | Rhodobacteraceae |
| bac51 | 0.82 | Bacteria | Cyanobacteria | Cyanobacteria (class) | Synechococcales | Synechococcaceae |
| bac99 | 0.48 | Bacteria | Bacteroidetes | Sphingobacteria | Sphingobacteriales | Saprospiraceae |
| bac127 | 0.40 | Bacteria | Bacteroidetes | Cytophagia | Cytophagales | Cyclobacteriaceae |
| bac151 | 0.33 | Bacteria | Bacteroidetes | Cytophagia | Cytophagales | Acidimicrobiaceae |
| bac240 | 0.21 | Bacteria | Actinobacteria | Acidimicrobiia | Acidimicrobiales |  |
| bac248 | 0.20 | Bacteria | Proteobacteria | Alphaproteobacteria | SAR11 clade |  |
| bac249 | 0.11 | Bacteria | Firmicutes | Clostridia | Clostridiales | Clostridiaceae |
| bac330 | 0.15 | Bacteria | Proteobacteria | Alphaproteobacteria | Rhizobiales | Rhodobiaceae |
| bac337 | 0.15 | Bacteria | Chloroflexi | Caldilineae | Caldilineales |  |
| euk3 | 11.1 | Eukaryota | Metazoa | Bryozoa | Gymnolaemata | Cheilostomatida |
| euk17 | 1.16 | Eukaryota | Stramenopiles | Bacillariophyta | Bacillariophyceae | Bacillariophycidae |
| euk18 | 1.63 | Eukaryota | Stramenopiles | PX clade | Phaeophyceae | Ectocarpales |

A partial Mantel test, correcting for all measured environmental variables, confirmed a strong correlation between bacterial and eukaryote community composition (partial mantel r = 0.92, p = 0.001), which was suggested by the similar clustering in the nMDS ordinations Fig. 2a-b. To further explore correlations between individual bacterial and eukaryotic sequence variants, a co-occurrence network was constructed (Fig. 5). The network features several prominent sequence variants that correlate with many other taxa and in some cases form distinct network modules. In particular, a sequence variant classified as *Hydrozoa (Metazoa: Cnidaria)* displayed both positive (co-occurrence) and negative (mutual exclusion) correlations with several bacterial taxa. Other prominent eukaryotic taxa were classified as *Mytilus* sp. (mussel), *Dorvillea* sp. (polychaete) and *Bacillariophyta* (diatoms). In some cases, network modules centered around Bacteria, such as *Bacteroidetes, Proteobacteria* and *Cyanobacteria*.

**Figure 5:**
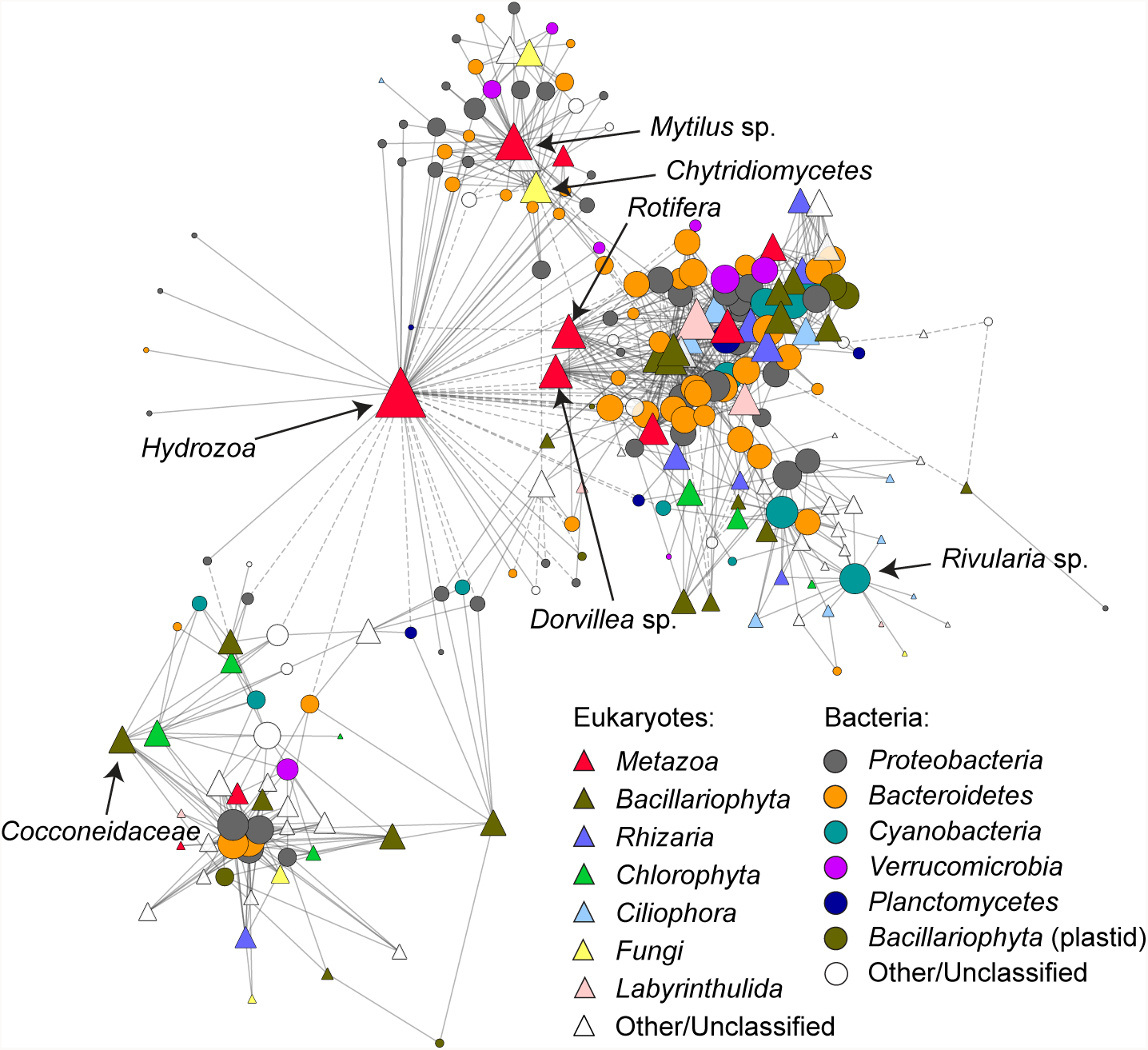
Co-occurrence of bacterial and eukaryotic rRNA sequence variants presented as a network. Solid lines (network edges) indicate significant positive Pearson correlations (r>0.8, p<0.01) between bacterial (circles) and eukaryotic (triangles) rRNA sequence variants (i.e. taxa, network nodes) while dashed lines indicate significant negative correlations (r>-0.8, p<0.01). Nodes are sized according to the number of correlating neighbors (degree) and color-coded according to taxonomic affiliation. Selected prominent nodes (same as in Fig. 4) are further labeled with more detailed taxonomic classifications.

### DISCUSSION

### Eelgrass bacterial microbiomes are locally variable

This study represents the first effort to characterize both bacterial and eukaryotic epibiotic communities on eelgrass leaves and to relate their communities to potentially important environmental drivers and to each other. We found that the local abiotic environment, in the form of water depth, explained a great deal of variation for both communities. Water depth was one of the main differences between lagoon and open coast sites which likely in part explains the clear separation observed between these sites. However, water depth encompasses separate physical factors such as light penetration and wave shear forces which are difficult to further separate in this study. In addition, several other factors such as mean salinity, nutrient levels and water temperature differ on seasonal time scales in lagoons compared to the open coast (Blindow et al., 2016;Schiewer, 2008) and could also contribute to this variation.

Of the biotic environmental variables measured, eelgrass shoot leaf area (a measure of the size of each shoot) and biofilm chlorophyll *a* content explained the most variation in community composition of both bacteria and eukaryotes. The explanatory power of eelgrass leaf area, which is related to eelgrass productivity (Echavarri &#x00ED; a-Heras et al., 2010), suggests an influence of host physiology on epibiotic biofilm composition. The amount of variation explained by chlorophyll *a* content, a proxy for the contribution of photosynthetic organisms (algae and cyanobacteria) in the biofilms, further indicates that interactions within the biofilms, for example between eukaryotic algae and bacteria play an important role in shaping community composition.

Richness of bacterial sequence variants was correlated with leaf area and with eelgrass dry weight at the sampling sites. Larger leaf area corresponds directly to a larger area of biofilm sampled, since biofilm from entire eelgrass shoots was removed. This observed richness-area relationship agrees with similar biodiversity-area relationships in other microbial habitats (reviewed in Martiny et al., 2006) and among macroscopic organisms. Eukaryotic biodiversity (measured as sequence variant richness) seemed to be controlled by factors specific to lagoon vs. open coast habitats, rather than area.

Despite the small area surveyed in this study, variability in community composition of both bacteria and eukaryotes was substantial among sites, which shows that eelgrass leaf biofilms are dynamic habitats. This observation is consistent with recent results from a global survey of eelgrass bacterial microbiomes that revealed that leaf surface microbiomes were more variable than root-associated microbiomes (Fahimipour et al., 2016). Nevertheless, we detected several sequence variants in all samples analyzed, supporting the possible existence of a core microbiome of eelgrass leaves. Several core sequence variants were classified to the family *Rhodobacteriaceae (Alphaproteobacteria),* which are often surface-attached, chemoorganotrophic bacteria common in the marine environment (Dang and Lovell, 2002). Interestingly, this family was also consistently detected in the rhizospheres of eelgrass and its close relative *Zostera noltii* (Cu &#x00FA;cio et al., 2016) as well as on leaves and below-ground parts of the seagrass *Halophila stipulacea* in the red sea (Mejia et al., 2016) indicating that their members indeed play important roles in association with various seagrasses. The observation of a *Methylophilaceae (Betaproteobacteria)* sequence variant as an abundant core community member agrees with detection of this taxon on eelgrass and other co-occurring macrophytes in the brackish-water Chesapeake Bay (Crump and Koch, 2008) and suggests importance of one-carbon metabolism on eelgrass leaf surfaces (Lapidus et al., 2011).

### Eelgrass bacterial microbiomes correlate with eukaryotic epibiont communties

The strong correlation between the bacterial and eukaryotic communities observed in this study has at least two possible causes. First, the abiotic and biotic environment experienced by both communities could influence them in similar ways. This explanation is partially supported by the similar amounts of variation explained by water depth, leaf area and chlorophyll *a* content for both communities. Second, bacterial and eukaryotic taxa may interact directly, resulting in a bacterial community that is driven by the composition of the eukaryotic community, or vice versa. The topology of the co-occurrence network suggests some importance of the latter scenario, where certain eukaryotic taxa (e.g. classified as *Hydrozoa, Mytilus* sp. and *Cocconeidaceae)* feature a disproportionate number of significant correlations with bacterial taxa. This indicates that bacterial communities in mature biofilms on eelgrass leaves are to some extent a reflection of the combined microbiomes of several epibiotic eukaryotes such as hydrozoans, mussels and diatoms in addition to any eelgrass-specific microbiome that may exist. Conversely, some bacterial taxa correlate with several eukaryotic taxa. For example, a cyanobacterial sequence variant classified as *Rivularia* sp. correlates with several ciliate taxa which may for example be attracted to this cyanobacterium as grazers.

## Outlook

To date, many studies addressing the microbiomes of marine organisms have focused exclusively on prokaryote communities (see e.g.Bourne et al., 2016;Hentschel et al., 2012; Wahl et al., 2012and references therein). In contrast, free-living microbial eukaryotes are receiving increasing attention and molecular community analysis has led to several recent discoveries that emphasize their underexplored diversity and crucial roles in the environment (Lima-Mendez et al., 2015;Liu et al., 2009). It is likely that insights of a similar caliber can be gained by focusing on microbial eukaryotes associated to macroscopic hosts (Andersen et al., 2013), beyond relatively well-described eukaryotic symbionts such as zooxanthella in corals (Muller-Parker et al., 2015). We have shown that the bacterial microbiome is strongly correlated with microbial eukaryote communities, indicating that interactions between eukaryotes and bacteria drive community assembly on eelgrass surfaces to some extent. By ignoring microbial eukaryotes in bacterial microbiome studies, a large part of the variation in bacterial communities could remain unexplained or misinterpreted. Indeed, the high diversity of both eukaryotic primary producers such as diatoms and heterotrophic taxa such as ciliates, rhizarians and many macroscopic and microscopic animals paint a picture of complex microbial ecosystems encompassing several trophic levels on eelgrass leaves. In these “microbial jungles”, bacteria are bound to play important roles which may be especially relevant in interaction with their microbial eukaryote neighbors.

In addition to considering the understudied microbial eukaryotes in microbiome studies, gaining a mechanistic understanding into microbiome assembly requires means to infer causative relationships between microbiome dynamics and abiotic and biotic factors (Widder et al., 2016). The present study included a limited number of samples from a small geographic area, providing a valuable first assessment of environmental drivers of community assembly and potential microbial interactions. Further insight could be gained by manipulative experiments, whereby for example, seagrass surfaces are exposed to colonization by microorganisms under controlled conditions. In addition, large observational microbiome datasets from a range of eelgrass meadows encompassing wider environmental gradients would allow more statistical power to tease apart different drivers of community assembly. Modeling approaches such as structural equation modeling (Pugesek et al., 2003) could allow assessment of causative relationships between the environment, the seagrass host and its microbiome. A major challenge lies in linking microbiome dynamics to processes that are relevant on the ecosystem level. Manipulative experiments that operate on an ecosystem scale, i.e. mesocosm experiments mimicking seagrass meadows, offer a promising approach to manipulate the environment while tracking microbiome dynamics and their links to critical ecosystem fluxes such as primary production and respiration. Ultimately, understanding assembly and function of seagrass microbiomes could contribute to tackling some of the challenges faced by seagrass meadow ecosystems under global climate change and other anthropogenic stressors.

## CONFLICT OF INTEREST STATEMENT

The authors declare no conflict of interest.

## AUTHORS AND CONTRIBUTORS

M.M.B. conceived the study, participated in the field work, processed the eelgrass samples, led the laboratory work, analyzed the data and wrote the paper. A. Bu&#x00FC; hler planned and executed the scuba diving, gathered the macrophyte data and prepared Fig. 1. A. Brauer performed molecular laboratory work and contributed to data analysis. S.D. participated in planning and execution of the field work and drove the research vessel. H.S. planned the fieldwork. I.B. planned and supervised the fieldwork and macrophyte data collection. All authors edited the manuscript.

## FUNDING

The field sampling campaign was funded through a grant from the German Federal Ministry of Education and Research (Project BACOSA, 03F0665C) awarded to H. S. and I. B. Microbial community analysis was funded through the University of Greifswald.

## ACKNOWLEDGEMENTS

The National Park administration (Nationalparkamt Vorpommern) kindly permitted us to dive and sample within the lagoon area. Tim Urich provided essential infrastructure and support for the microbial community analysis.

